# ASSESSMENT OF ALPHAFOLD PROTEIN MODELS FOR SMALL-MOLECULE LIGAND DOCKING

**DOI:** 10.64898/2026.01.04.697577

**Authors:** Hlib Maidanik, Maria Lazou, Rohin Bajaj, Raheel Sarwar, Sandor Vajda, Diane Joseph-McCarthy

## Abstract

Molecular docking is a powerful computational tool for predicting protein-ligand interactions, widely employed in drug discovery. However, its effectiveness is often constrained by the availability of experimentally resolved X-ray protein structures, a process that is both time consuming and resource-intensive. AlphaFold (AF), a deep learning method, offers an efficient alternative by predicting high-accuracy 3D protein structures directly from amino acid sequences. This study assesses the utility of AF-generated protein models for fragment and larger ligand docking with Glide, a widely used docking approach. The docking workflow is evaluated in an unbiased manner by carrying out binding site identification with FTMap, a binding hot spot prediction software. We show that fragment docking to AF models outperforms docking to the respective unbound protein crystal structures, and performs comparably to docking to the corresponding ligand bound structures when using an unbiased approach. Leveraging computational efficiency of AF model generation, we also employ ensembles of AF models to incorporate protein flexibility. Results show that docking to AF ensembles improves larger-ligand docking compared to docking to singular AF models and outperforms docking to unbound structures. The results provide insights into the effectiveness of integrating AF protein models into docking procedures, highlighting the potential for streamlining computational drug discovery processes.

**STATEMENT OF SIGNIFICANCE:** This work addresses a critical bottleneck in computational drug discovery by demonstrating that AlphaFold (AF) models can serve as an alternative or complement to experimental structures for molecular docking. Specifically, a systematic study assessing Glide docking to rigid AF protein models and ensembles of models compared to experimentally determined ligand-bound and unbound protein X-ray structures was performed. The evaluation employs an unbiased methodology using FTMap-identified binding sites, eliminating the need for prior knowledge of the native ligand binding location. Additionally, protein flexibility is incorporated through a multiseed ensemble approach that generates a conformational ensembles of AF models at minimal computational cost, improving the docking accuracy without the need for ligand-bound templates.

## INTRODUCTION

Molecular docking is a computational structure-based drug discovery technique that predicts the binding poses and relative affinities of a small-molecule ligands within target protein binding sites. This approach enables rapid screening of vast compound libraries, accelerating hit-to-lead optimization while reducing experimental cost and providing mechanistic insight into protein-ligand interactions (1). However, molecular docking functionality is inherently constrained by the availability of high-resolution experimental protein structures, which require time-intensive and resource demanding X-ray crystallography, Nuclear Magnetic Resonance spectroscopy, or cryo-Electron Microscopy. The recent advent of AlphaFold (AF), a neural network-based protein structure prediction model, has significantly expanded the availability of accurate protein structures (2–4). As such, AF offered unprecedented potential to overcome the experimental structure bottleneck that has historically limited structure-based drug discovery methods. While AlphaFold2 (AF2) models have previously been assessed for use in molecular docking with a variety approaches, the results have been mixed (5–10).

AF is trained predominantly on well-folded, stable PDB structures whereas active sites, which are inherently flexible to enable catalytic function, do not necessarily follow the same dominant principles of protein folding, making them more difficult to predict (5). Moreover, AF’s training set is heavily biased toward thermodynamically stable inactive conformations, potentially overlooking transient active states critical for drug binding. Scardino et al. conducted a benchmarking study using four docking programs (AutoDock, PLANTS, rDock, ICM) on well-characterized targets, and demonstrated that docking to AF models performed worse than docking to experimental holo structures (6). Despite overall backbone accuracy, small variations in side chain orientations had a disproportionate impact on docking outcomes. Karelina et al. showed that while AF2 captured the position of binding site residues more accurately than homology models for G-protein coupled receptors (an important but difficult class of proteins to model), docking performance was not improved relative to homology models. Holcomb et al. also found a performance gap between using AF2 models and holo structures for docking over the PDBbind dataset using AutoDock-GPU (8). They also showed that docking to AF2 models performed comparably to docking to apo structures suggesting that predicted models may capture ligand-bound conformations to some degree. Zhang et al. demonstrated that refinement of the AF2 models using an aligned known binding ligand as the template improved the docking, and that Glide-generated docking poses of known binders could also similarly be used as templates for IFD-MD (9). This refinement while effective introduced bias into the docking in that it required a known ligand binder which does not always exist. Diaz-Rovira et al. developed an AF2 with templates with more than 30% identity to their test set excluded in an attempt to mimic a more real world virtual screening situation (10).

Despite these findings, critical gaps remain in understanding how to utilize AF models effectively for structure-based virtual screening campaigns. In this study, we expand upon the prior work in several important ways: (i) our test set consists of fragment and corresponding larger-ligand bound structures as well as unbound structures, allowing us to evaluate the utility of AF2 to dock different sized small molecules; (ii) we compare Glide XP and SP; (iii) we evaluate docking fragments to AF2 ensembles generated using a multi-seed approach with templates removed entirely. In addition, we describe a novel protocol for large-scale structure-based virtual screening when no structure of the protein or known ligand binders exist. The approach combines FTMap (11,12) for binding site identification with Glide docking (13–15) to AF2 ensembles.

Accounting for protein flexibility remains a persistent challenge in molecular docking in general (16,17). Proteins undergo conformational rearrangement upon ligand binding, yet most docking algorithm, including the Glide software (13,14), treat receptors as rigid entities (18,19). The stochastic nature of AF model generation (2) can be leveraged to incorporate protein flexibility at minimal computational cost. By intentionally varying the random seed used during multiple sequence alignment (MSA) generation across independent AF runs, alterations in MSA depth and composition produce conformationally heterogeneous ensembles of protein models (20–22). The multiseed AF ensemble approach represents a template-free methodology for sampling protein flexibility.

This study systematically evaluates the utility of AF2 models for fragment and larger lead-like ligand docking with Glide (13–15), comparing performance against experimentally resolved ligand-bound and unbound X-ray structures. We employ an unbiased methodology utilizing FTMap computational hot spot mapping (11,12) to identify target binding sites for docking. We show that fragment docking to AF models performs similarly to experimental unbound structures while AF ensembles can improve docking accuracy for larger ligands, surpassing that of unbound structures. A workflow for early-stage drug discovery when experimental structures and known small-molecule binders are unavailable is described.

## METHODS

### Protein Test Set

For this study, a previously published benchmark set which consists of 33 proteins each with at least one corresponding fragment bound, larger ligand bound, and unbound X-ray structure was used (23). For each protein in the test set (see **Table 1**), the sequence from the fragment-bound structure was used to retrieve the corresponding model from the AlphaFold Protein Structure Database (EMBL-EBI) (2, 24). Each fragment-bound X-ray structure was taken as the reference to which the respective larger-ligand bound structure, unbound structure, and AF protein model were aligned using PyMol (The PyMOL Molecular Graphics System, Version 3.1.5.1, Schrödinger, LLC).

**TABLE 1.**
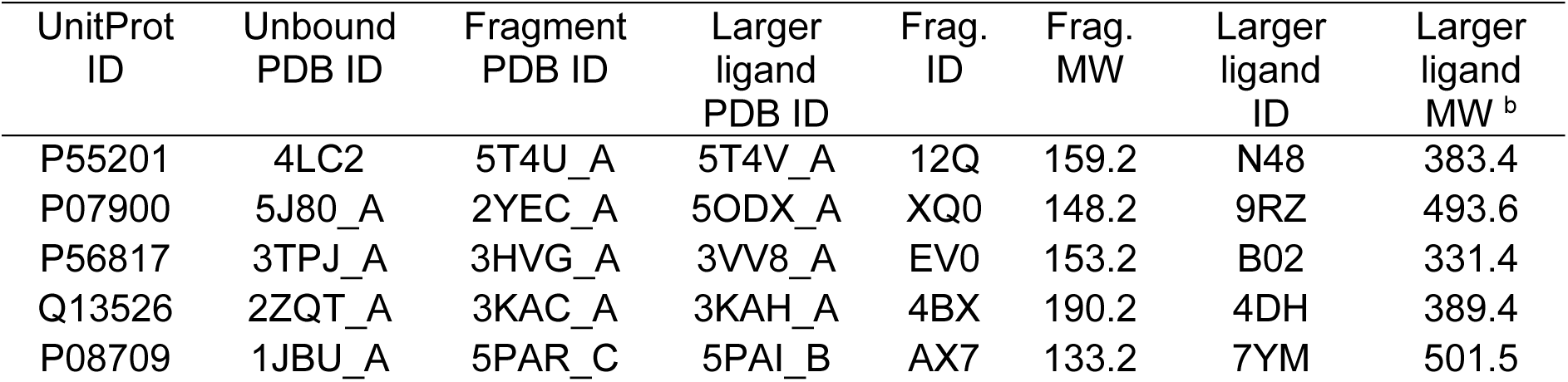
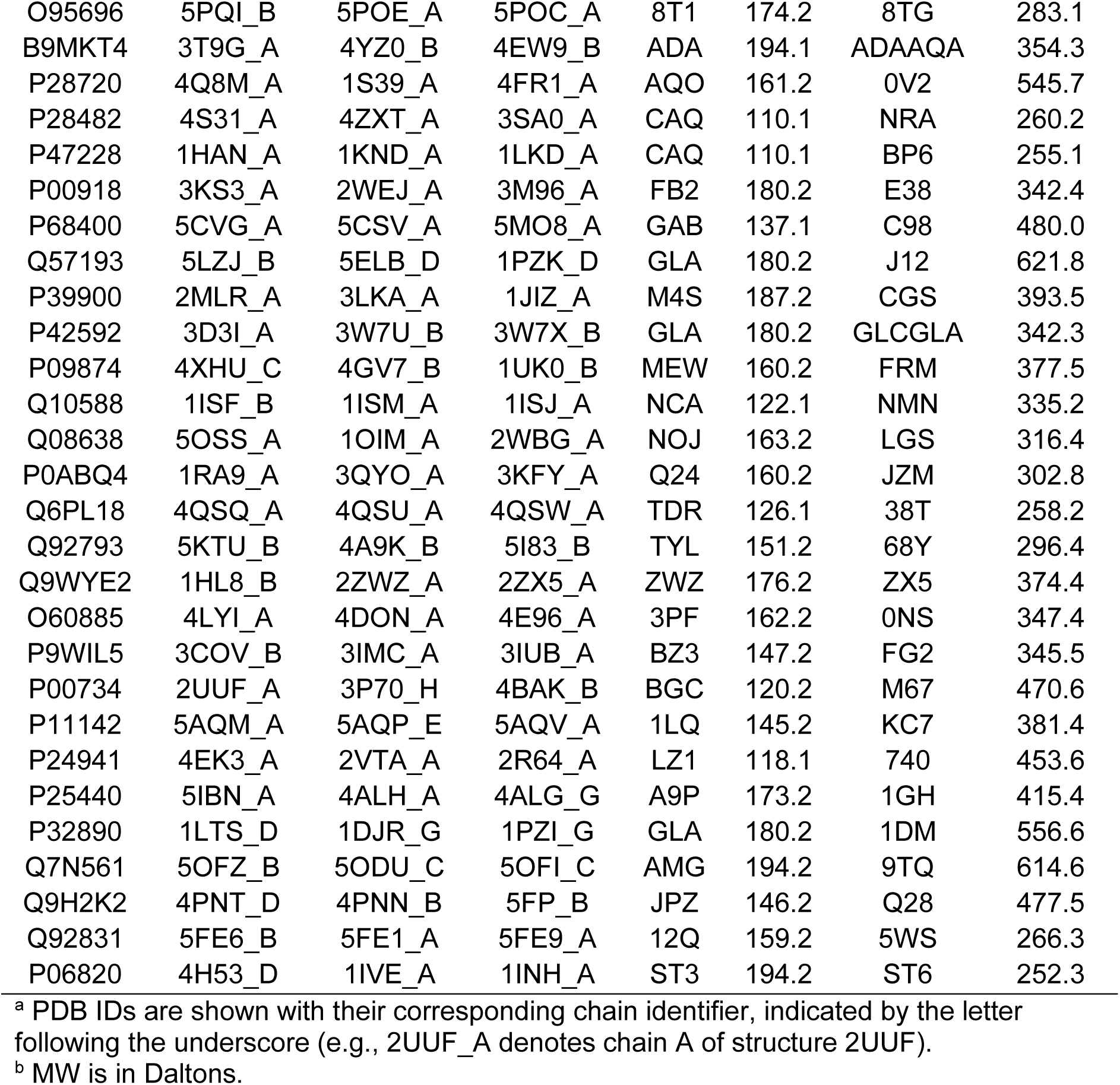
Protein test set ^a^.

### Generation of AF Ensembles

Ensembles of AF models were generated using ColabFold (21,22), an open-source implementation of the AF2 algorithm. The MSA inputs were generated using MMseq2 (25). The number of initial seeds for MSA initialization during inference stage in ensemble generation workflow was set to 100, and AF outputs 5 models per seed, resulting in 500 structural models per protein sequence. For a given protein, the top 100 models were selected as the “AF ensemble” based on the mean per-residue predicted local distance difference test (pLDDT) score (2).

### Hot Spot Mapping to Define Docking Box

To facilitate unbiased docking, binding site hot spots were calculated using FTMap (11,12) and top scoring hot spots were used to define the binding site box for docking. Specifically, the set of unbound structures and AF-generated protein models were subjected to binding hot spot mapping using the FTMap server. FTMap locates binding site hot spots on the surface of proteins, through exhaustive sampling of the position of 16 small organic probe molecules on the protein interface. The number of probe clusters in each hot spot establishes its rank, with the top-ranked hot spot containing the highest number of probe clusters. Kozakov et al. set a druggability criterion for hot spots such that those that contain 13 to 15 probe clusters are borderline druggable, while hot spots with 16 or more may be druggable (26). Based on this, only hot spots with 13 or more probe clusters were selected to define the binding site box for docking. For AF Ensembles, the mapping results for the corresponding AF Model were used to define the box.

### Protein and ligand preparation

The aligned protein structures, models, were prepared using default parameters. This preparation included the assignment of protonation states, optimization of hydrogen bonds assignments, enumerate bond orders to HET groups, and minimal energy minimization (27). Ligands (fragments and larger ligands) were prepared using the LigPrep tool in Maestro (Schrödinger Release 2024-2: Schrödinger, LLC, New York, NY, 2024), which optimizes ligand geometries and assigns tautomeric and ionization states with default parameters.

### Docking Grid Generation

Receptor grids for Glide docking (13, 14) were generated in Maestro. The computed grids represent the energy landscape of the target protein binding site as a set of force fields, allowing for more accurate and efficient energy evaluation during docking. The grid is defined in 3-dimensional space by concentric inner and outer boxes: the inner box specifies the allowed positions for the site-point search, while the outer box constrains the space in which all ligand atoms must be contained. The workflow requires a centroid around which the boxes are generate; we defined this centroid using a workspace ligand. The size of the inner box was set to the default 10×10×10 Å^3^; however, the outer box size setting varied as described below, depending on the ligands used to define the box center.

We used either a native X-ray bound ligand or a central probe from a top-ranked FTMap consensus site for grid generation for a given protein. By default, the outer box size is set based on the assumption that similarly sized ligands will be docked. Alternatively, the box size can be manually specified with the default dock ligand of size ≤ 20 Å (resulting in an overall box size of 30×30×30 Å^3^; see **Figure S1**). We performed docking to all sets of protein structures and models using three different combinations of box center and size configurations: (1) bound ligand center with the box size automatically determined based on workspace ligand size, (2) bound ligand center with dock ligand of size ≤ 20 Å, and (3) FTMap probe center with dock ligand of size ≤ 20 Å (as summarized in **Table 2**). Configuration (3) defines the binding site for docking in an unbiased and ligand-agnostic manner.

**Table 2.** Description of Box Configurations.

|  | **Box Center** | **Enclosing Box Size Setting** |
| --- | --- | --- |
| **1** | Centroid of bound ligand | Dock ligands similar in size to ligand |
| **2** | Centroid of bound ligand | Dock ligands with length ≤ 20 Å |
| **3** | Centroid of central hot spot probe | Dock ligands with length ≤ 20 Å |

### Reproducing the X-ray Ligand-Protein Complexes

The Glide (13, 14) version 10.2.138 in Maestro (Schrödinger, LLC, New York, NY, 2024) was utilized for docking ligands to the pre-computed receptor grids. The docking settings were adjusted for XP and saving up to five poses per ligand (15). As a control, each ligand was docked to its corresponding native bound protein structure. This was accomplished by docking the respective ligands to receptor grid generated based on the centroid of the bound ligand in the X-ray structure and the box size selected automatically for docking similarly sized ligands (box configuration 1). This method provides a baseline for the accuracy of the docking protocol by evaluating the results under the ideal condition. For a comprehensive comparison to docking to the unbound protein structures and AF models, the fragments and larger ligands were also docked to their corresponding bound protein structures with box configurations 2 and 3. Glide SP docking was also investigated where specified.

### Fragment Docking

Each fragment was subsequently docked into the corresponding unbound protein structure (see **Table 1**), AF model, and AF ensemble using utilizing the receptor grid generated using a central probe from the top ranked hot spot. However, if the best RMSD out of the top 5 poses was greater than 2 Å, the ligands were docked to the grid generated by the next best ranked hot spot. This process was repeated until a pose with RMSD under 2 Å was found or there were no additional hot spots with a score ≥ 13.

### Larger Ligand Docking

For each fragment bound protein structure there is a corresponding larger ligand bound structure. Those larger ligands contain the corresponding fragments in their structure. Superimposing fragment and larger ligand bound PDBs shows that the fragments align well with the corresponding larger ligand parts. After evaluating the accuracy of small fragment docking, larger ligands were also docked. The larger ligands were docked to the receptor grids generated with the same central cluster probe in the site that performed best when docking the fragments. The docking settings were also adjusted to XP, saving up to 5 poses.

### Ensemble Docking

For each protein in the test set, Glide docking of larger ligands was performed on the AF ensemble of models generated via multiseed approach. Custom Python and bash scripts were written and implemented to automate protein model alignment with the reference X-ray structure, minimization, receptor grid generation, and docking. Previously prepared ligands were used for docking.

### Induced-Fit Docking

Induced-fit docking (IFD) protocol (Schrödinger, LLC, New York, NY, 2025) was employed to dock larger ligands to AF models (28, 29). IFD module is available through Maestro interface, hence previously prepared AF models and larger ligands were used. IFD can use either Glide SP or Glide XP (after the initial placement with Glide SP). Herein Glide XP was specified using Schrödinger Release 2024-2 and Glide SP with Schrödinger Release 2025-3. Receptor grids were rendered using box configuration 1. Ranking of poses was done using the standard IFD Score metric. The output for each protein system was a protein conformation with five ranked poses for the ligand, followed by the next best protein conformation (in terms of the best IFD Score) with its five ranked poses for the ligand, and so on. So, the Top 5 poses were for the top ranked complex and the Top 10 poses were for the top two complexes.

### Pose Analysis

As part of the designed docking pipeline, all docked poses and their corresponding docking scores were exported using Schrödinger’s *structconvert* utility command. The heavy atom root-mean-square deviation (RMSD) between the docked ligand pose and the corresponding X-ray bound ligand position were computed to assess the accuracy of the docking. The generated docked poses were exported as separate files together with the experimental bound ligand from the structure that was previously aligned in PyMol. Schrödinger’s script *rmsd.py* was then used to calculate RMSD between each pose and their respective reference bound ligand.

## RESULTS and DISCUSSION

### Reproducing Protein-Ligand Complexes

The docking of each fragment (**Table 3**) and larger ligand (**Table 4**) to its corresponding bound structure provides an upper limit on the docking accuracy. For box configuration 1, with ligand-biased center and size, fragments docked successfully with RMSDs ≤ 2 Å (≤ 2.5 Å) for 24% (33)% of the cases for the top pose, and 49 (61)% of the time if the top 5 poses were considered. We see a slight increase in fragment docking performance for the top pose with box configuration 2 (ligand-biased center and constant box size), with RMSDs ≤ 2 Å (≤ 2.5 Å) in 33 (42)%. The improvement of results from configuration 1 to 2 suggests that slightly increasing the search space dimensions improves the fragment docking. Box configuration 3, the unbiased case, yields results similar to box 1 for the top ranked pose with RMSDs ≤ 2 Å (≤ 2.5 Å) in 21% (39%) of structures, indicating that utilizing a central probe from a strong hot spot to define the box center is an effective alternative to utilizing the bound ligand. These success rates for fragment are consistent with previously reported values of similar docking experiments (30, 31).

**TABLE 3.** Fragment docking performance across the test set.

| Box Configuration | Pose Rank<sup>a</sup> | RMSD $\leq 2$ Å | | | RMSD $\leq 2.5$ Å | | |
| --- | --- | --- | --- | --- | --- | --- | --- |
|  |  | X-ray Bound [%] | X-ray Unbound [%] | AF Model [%] | X-ray Bound [%] | X-ray Unbound [%] | AF Model [%] |
| 1 | Top 1 | 24.2 | 21.2 | 24.2 | 33.3 | 24.2 | 39.4 |
|  | Top 5 | 48.5 | 39.4 | 39.4 | 60.6 | 48.5 | 57.6 |
| 2 | Top 1 | 33.3 | 15.2 | 18.2 | 42.4 | 18.2 | 24.2 |
|  | Top 5 | 54.4 | 39.4 | 42.4 | 63.6 | 45.5 | 54.5 |
| 3 | Top 1 | 21.2 | 21.2 | 33.3 | 39.4 | 30.3 | 36.4 |
|  | Top 5 | 45.5 | 36.4 | 54.5 | 51.5 | 45.5 | 57.6 |

<sup>a</sup> Results are for the top ranked pose and the lowest RMSD pose among top 5 poses

**TABLE 4.** Larger-ligand docking performance across the test set.

| Box Config. | Pose Rank<sup>a</sup> | RMSD $\leq 2$ Å | | | | RMSD $\leq 2.5$ Å | | | |
| --- | --- | --- | --- | --- | --- | --- | --- | --- | --- |
|  |  | X-ray Bound [%] | X-ray Unbd. [%] | AF Model [%] | AF Ens. [%] | X-ray Bound [%] | X-ray Unbd. [%] | AF Model [%] | AF Ens. [%] |
| 1 | Top 1 | 21.2 | 21.2 | 12.2 | 18.2 | 27.3 | 24.2 | 18.2 | 24.2 |
|  | Top 5 | 33.3 | 30.3 | 24.2 | 33.3 | 39.4 | 30.3 | 30.3 | 33.3 |
| 2 | Top 1 | 24.2 | 18.2 | 21.2 | 18.2 | 30.3 | 21.2 | 21.2 | 27.3 |
|  | Top 5 | 45.5 | 30.3 | 27.3 | 30.3 | 51.5 | 30.3 | 27.3 | 33.3 |
| 3 | Top 1 | 18.2 | 15.2 | 12.2 | 18.2 | 24.2 | 18.2 | 18.2 | 27.3 |
|  | Top 5 | 39.4 | 24.2 | 18.2 | 30.3 | 45.5 | 24.2 | 24.2 | 33.3 |

<sup>a</sup> Results are for the top ranked pose and the lowest RMSD pose among top 5 poses

For the larger ligand docking, the accuracy is comparable to that for fragment docking when considering the top pose with RMSDs ≤ 2 Å for all box configurations. However, the performance is generally poorer among the top 5 docked poses with 39% vs. 46% (33% vs. 39%) of the proteins having a pose ≤ 2 Å when using box configuration 3 (box configuration 1). Poses with RMSDs ≤ 2.5 Å in top 5 were found for 46% vs 52% (39% vs. 61%) of cases using box configuration 3 (box configurations 1). These results are also consistent with published findings demonstrating that docking accuracy declines as ligand size increases due to a higher number of rotatable bonds present (32). The mean number of rotatable bonds in the fragments in our dataset was 0.7 (SD= 0.7, range= 0-2), compared to 5.2 for the much more flexible larger ligands (SD= 3.3, range= 0-14).

For box configuration 3 (unbiased), FTMap hot spots were used to define the binding site box as described in the Methods. For both the unbound structures and the AF models, a hot spot was identified near the correct ligand binding site for approximately 91% (30/33) of proteins (see **Table S1**). When hot spot mapping did not identify the correct binding site, the docking obviously was not success; this accounted for about 9% of the failures.

### Fragment Docking to the Unbound Structure vs. AF Model

The assessment of fragment docking to the AF model vs. the unbound structure is also summarized in **Table 3**. As displayed in **Figure 1**, the box used for the unbiased docking grid generation for these results were mainly derived from the top-ranked FTMap consensus site, site 0, although there were cases where consensus sites 1, 2 or 3 were used. For docking to the AF models with the unbiased box configuration 3, 33% of the cases yielded RMSDs ≤ 2Å in the top pose and 55% in the top 5 poses; this is compared to 21% (top 1) and 36% (top 5) when docking to the unbound structure. Overall, the accuracy of fragment docking to AF model is substantially higher compared to that for docking to the corresponding experimental unbound structure, especially when near correct poses are considered, and are better or similar to the results obtained when docking to the corresponding bound structure. These results agree with our previously published work showing that AF2 models are sufficient for hot spot mapping (33). They also suggest that AF2 may generate binding site geometries that are more consistent with experimentally determined bound structures compared to unbound structures which likely stems from the presence of more holo structures its training set (34).

**FIGURE 1.**
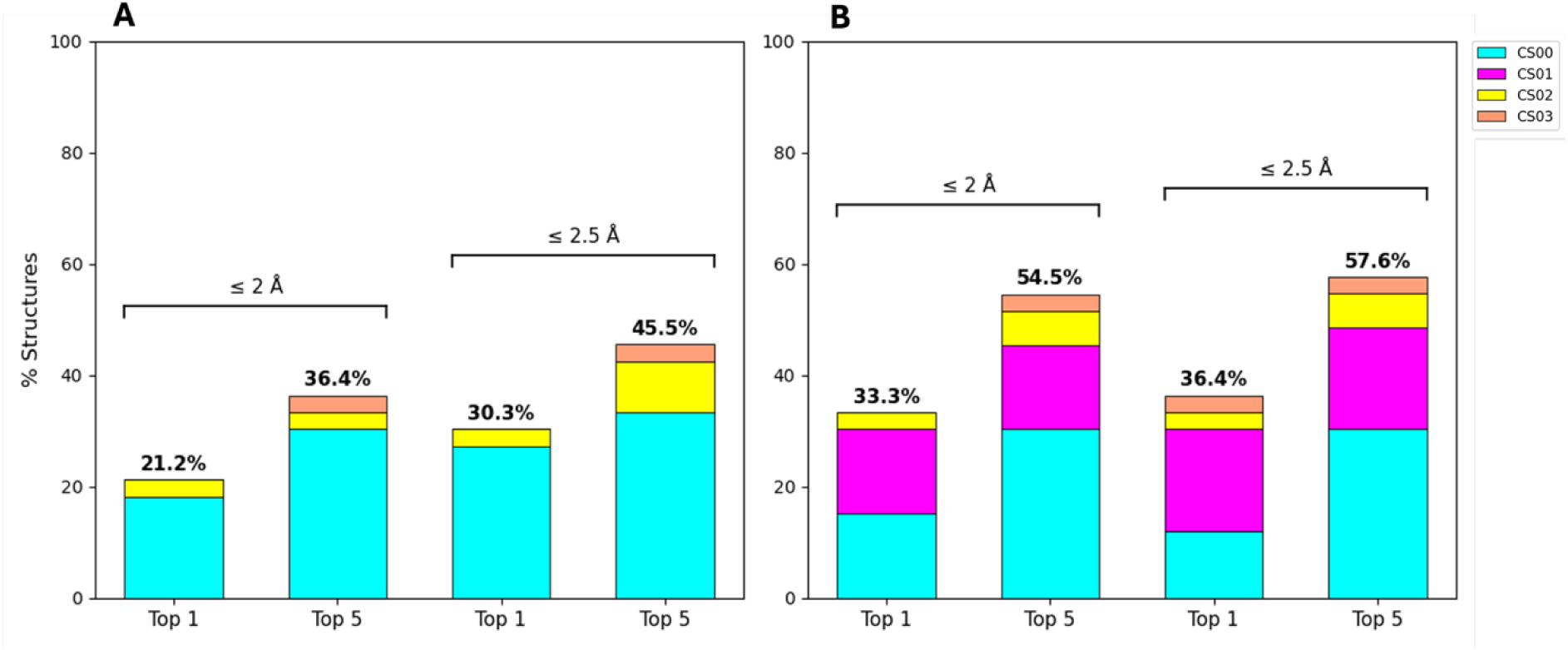
Comparison of correct (RMSD ≤ 2 Å) and near correct (RMSD ≤ 2.5 Å) fragment poses across the protein test set. Results are shown in (A) for unbound structures and (B) AF models for the top ranked pose and the best-RMSD pose among top 5 ranked poses. The color of the bars indicates the FTMap consensus site (CS) used for the docking box generation as indicated by the legend, where CS00 is the top ranked FTMap site, CS01 is the next highest ranked consensus site, and so on.

Moreover, using box configuration 3 (with the unbiased box center and size) the docking accuracy to AF models is generally improved or equivalent compared to that for when using the other box configurations. For example, the top pose has an RMSD ≤ 2 Å for 33% vs 18% vs 24% of proteins for box configurations 3 vs. 2 vs. 1, respectively. Similarly, the top pose has a near-correct pose (with an RMSD ≤ 2.5 Å) for 36% vs 24% vs 39% of proteins. This result is consistent with the fact that others have found that using receptor grid dimensions which are ∼3 times larger than the radius of ligand gyration leads to higher docking accuracy (35). It also indicates that using bound ligand coordinate centroid as box center captures the intended search space less effectively than using the coordinates of the computationally determine binding hot spot.

The RMSDs per protein for the top pose and best RMSD pose of the top 5 are shown in **Figure 2**. Sometimes the top ranked FTMap consensus site (Site 0) is not the site where the fragment binds which necessarily results in very large RMSDs (left plots in **Figure 2**). When the FTMap consensus site (with a score of ≥ 13) that gives the best RMSD to the X-ray ligand is used (middle plots in **Figure 2**), the mean RMSDs are therefore reduced. As anticipated when considering the top 5 poses using the best FTMap consensus site (right plots in **Figure 2**), the mean RMSD is further reduced for docking to the AF models and unbound structures. The mean RMSD across the test set drops from 4.8 to 3.8 Å when docking to the unbound structures, and from 4.1 to 3.0 Å when docking to the AF models. If the top 5 poses are considered, both mean RMSDs decrease as expected but that for the AF models is lower than that for the unbound structures. When considering the top 5 poses, the majority of the time a near native pose for the ligand is found when using the AF models. These findings indicate that accurate fragment docking is enabled using AF models with an unbiased box definition.

**FIGURE 2.**
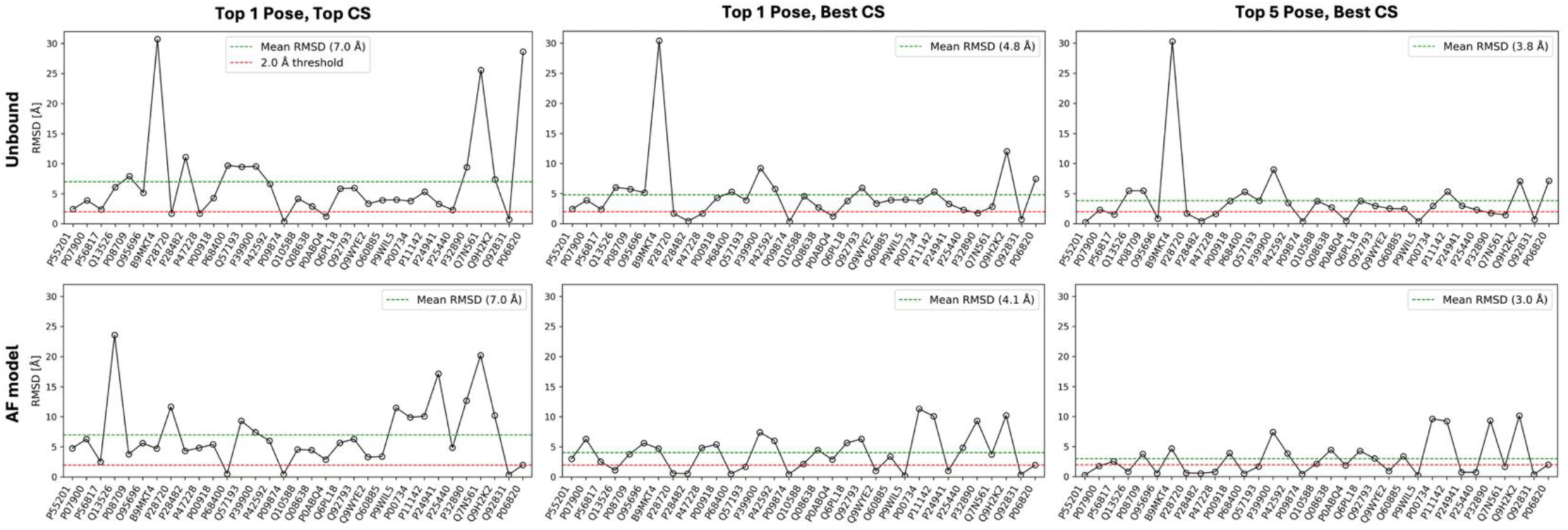
Fragment docking RMSDs per protein. The top row shows the results for docking to the AF model and the bottom row shows that for docking to the unbound structure. For each row, the left graph shows the RMSDs of the top pose for the docking box generated at Site 0, the middle graph shows the RMSDs of the top pose at the best consensus site, and right graph shows the best RMSD of top 5 poses at the best consensus site. The green line represents the mean RMSD across the test set, while the red line indicates the 2 Å cutoff for correct poses.

### Larger Ligand Docking to Unbound Structure vs. AF Model vs. AF Ensemble

The results for docking the larger ligands (with molecular weights ranging from 252 to 621 Da), to the X-ray structures vs. AF models and AF ensembles are summarized in **Table 4 and Figure 3**. The results for docking the larger ligands are similar to those for the fragment docking with box configuration 3 (unbiased). As anticipated, however, the docking accuracy decreases somewhat for the larger ligands with more rotatable bonds vs. the fragments. For the larger ligands, correct poses are obtained for 18% of the proteins when docking to the bound structure, and 15% when docking to the unbound structure. While for the fragments the results are improved for docking to the AF model relative to the unbound structure, the opposite is true for the larger ligands. For the larger ligands, the success rate is 12% for docking to the AF model compared to 15% for docking to the unbound structure. The docking success rate is similar for the biased vs. unbiased docking box configurations for both AF models and unbound structures.

**FIGURE 3.**
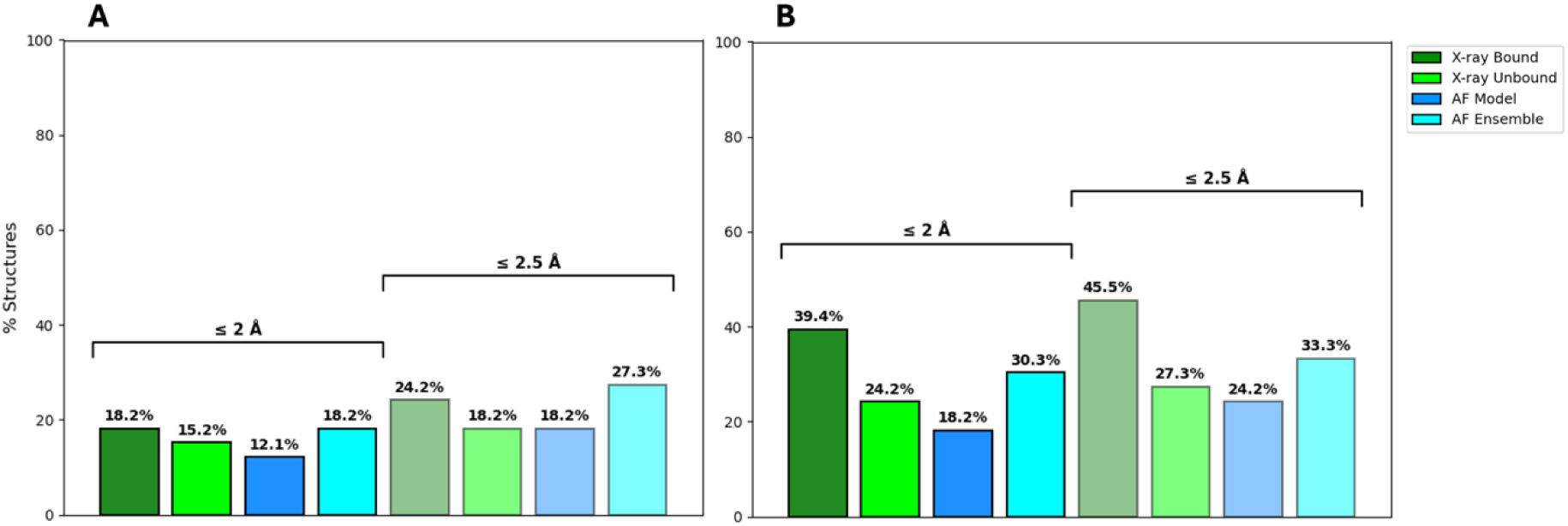
Comparison of correct (RMSD ≤ 2 Å) and near correct (RMSD ≤ 2.5 Å) larger-ligand poses across the protein test set. Results are shown in (A) for the top ranked pose and in (B) for the best-RMSD pose among top 5 ranked poses.

In an attempt to address the small decrease in performance with the AF models vs. the unbound structures for the larger ligands, ensembles of structures were generated for each protein using AF with multiple seeds as described in the Methods. Docking to the ensemble increases the performances from 12% (AF model) to 18% which is greater than the 15% for the unbound structure. A similar improvement is observed for top 5 as for top 1 and for correct and near correct poses.

IFD docking to the AF models was also attempted. Examining near correct poses (≤ 2.5 Å) shows that the standard Glide XP docking to the AF ensembles significantly outperforms the IFD docking for the top 1 pose (24.2 % vs. 18.2 %). For the top 5 and top 10 poses, the performance is similar (see **Figure S2**). Similar results were obtained using IFD with Glide SP (**Figure S3**).

The RMSDs per protein for docking the larger ligands to the unbound structures vs. AF models vs. the AF ensembles are shown in **Figure 4**. Docking to the unbound structures and AF models yields comparable results for the top ranked pose with mean RMSD values of 6.5 Å for both. Using AF ensembles improves docking outcomes compared to AF models for the top ranked pose as shown by the decrease in average RMSD value from 6.5 Å to 5.6 Å. Expanding the consideration to top 5 poses, we see a drop in the mean RMSD values for the three sets. While for the top 5 poses the unbound structures performing slightly better than AF models (5.5 vs. 5.8 Å), AF ensembles outperform both with a mean RMSD of 5.0 Å. The RMSD for docking to B9MKT4 unbound structure is very high (∼30 Å) as is the RMSD for docking to the Q9H2K2 AF model (∼23 Å); in both cases the high RMSD is due to the fact that FTMap did not identify the binding site with a hot spot scoring ≥ 13. For Q9H2K2 the binding site is identified with the unbound structure but the RMSD is still quite high (7.5 Å) (see **Table S2**). For P68400, FTMap correctly identified the active site with high-scoring hot spots in both the unbound structure and AF model, yet both yielded high RMSD values. This outcome likely stems from the large size (∼480 Da) and high flexibility (11 rotatable bonds) of the docked ligand. More detailed explanations of the high RMSD values observed in Figure 4 for the unbound structures vs. AF models are also provided in **Table S2**. Docking to the AF Ensembles lowers the RMSD of the top pose below 2Å for two proteins—P00918, and Q9WYE2—compared to docking to the AF models. For the top 5, it improves the RMSD to below 2 Å relative to docking to the AF models for five models—P55201, P07900, P28720 P00918, and P24941. Overall unbiased docking to the AF Ensemble outperforms docking to the unbound structures and AF models.

**FIGURE 4.**
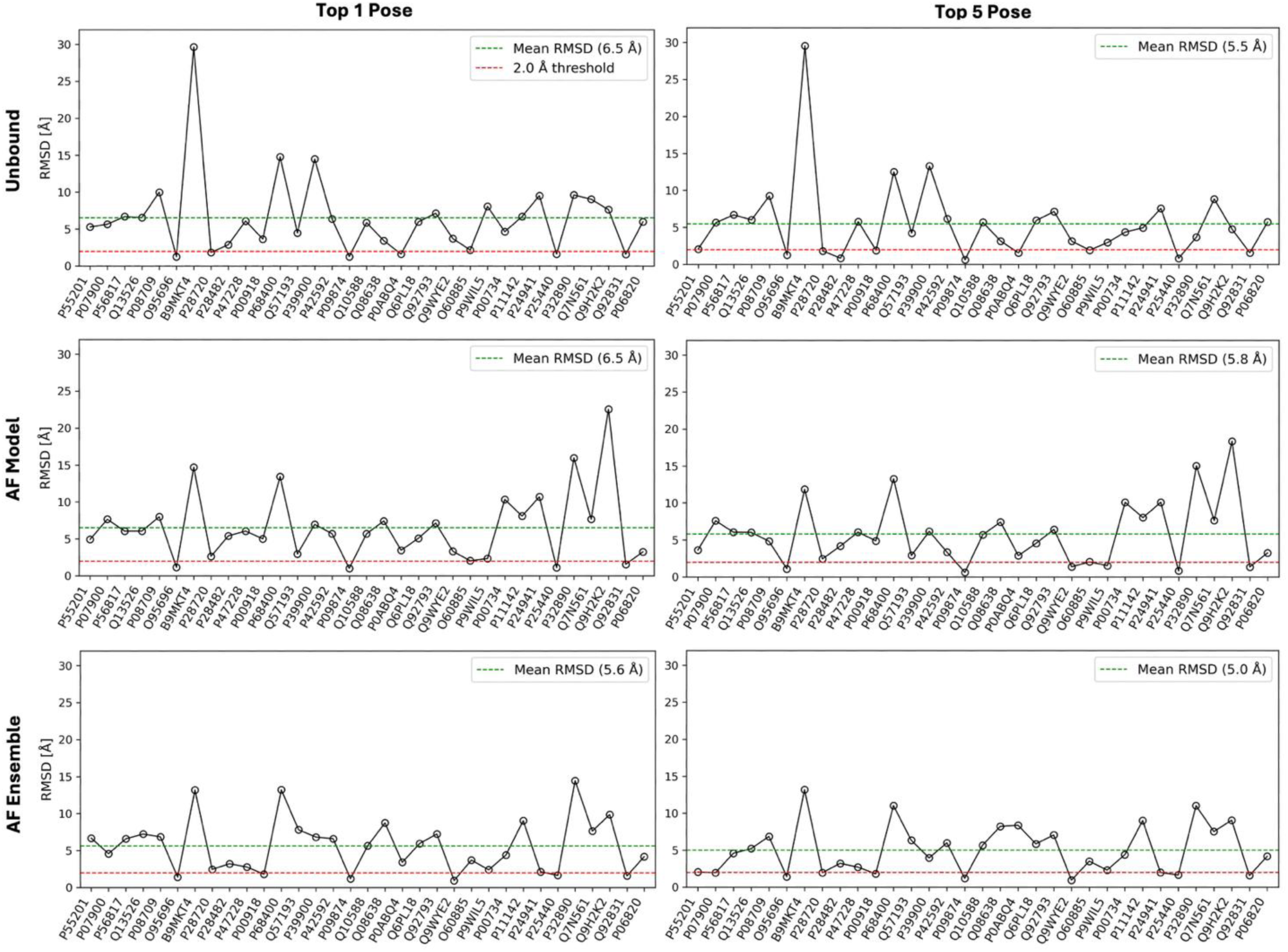
Larger-ligand docking RMSDs per protein. The top row shows the results for docking to the unbound structures, the middle row shows that for docking to the AF models and the bottom row for AF Ensembles. For each row, the left graph shows the RMSDs of the top pose, and the right graph shows the best RMSD of top 5 poses. The green line represents the mean RMSD across the test set, while the red line indicates the 2 Å cutoff for correct poses.

Ligand docking was also attempted with Glide SP. For fragments the results were similar to Glide XP (**Table S3**). However, for the larger ligands, the accuracy of docking to the AF models and AF ensembles degraded (**Table S4**). The enhance sampling and more intricate scoring function of Glide XP appears to be necessary for docking to the AF models in general.

### AF Ensemble Heterogeneity and Docking Accuracy

To explore the incorporation of protein flexibility through the generation of ensembles of AF models further, we considered two ensemble-selection approaches for capturing conformational heterogeneity: (i) the top 100 AF models ranked by pLDDT score, representing the most confident predictions across the full sampling space (used for the docking experiments described above), and (ii) the single highest-ranked pLDDT model from each of 100 random seeds to potentially further enhance structural variety via seed-based sampling. **Figure 5** illustrates that the two approaches produce nearly indistinguishable global conformational distributions (**Fig.5A** vs. **Fig.5B**) and marginally greater differences at the binding sites (**Fig.5C** vs. **Fig.5D**) for a subset of ensembles. Pairwise whole-protein RMSD heatmaps for the top 100 model ensembles (**Fig. 5A**) reveal that 19 out of 33 ensembles exhibit minimal conformational disparity (predominantly dark to light green, 0 to 1.5 Å), whereas 14 ensembles exhibit moderate to substantial variation (ranging from light green to dark red, 1.5 to 4 Å). In contrast, the conformational heterogeneity of the binding sites (**Fig. 5C**) is much smaller: 28 ensembles display nearly uniform dark green regions, indicating <0.5 Å RMSD variation, and only 5 ensembles display as light green to dark red. In addition, the mean pLDDT for the top 100 model ensemble 93.6 compared to 93.5 for the best model per seed; the mean binding site pLDDT was 95.0 for both.

**FIGURE 5.**
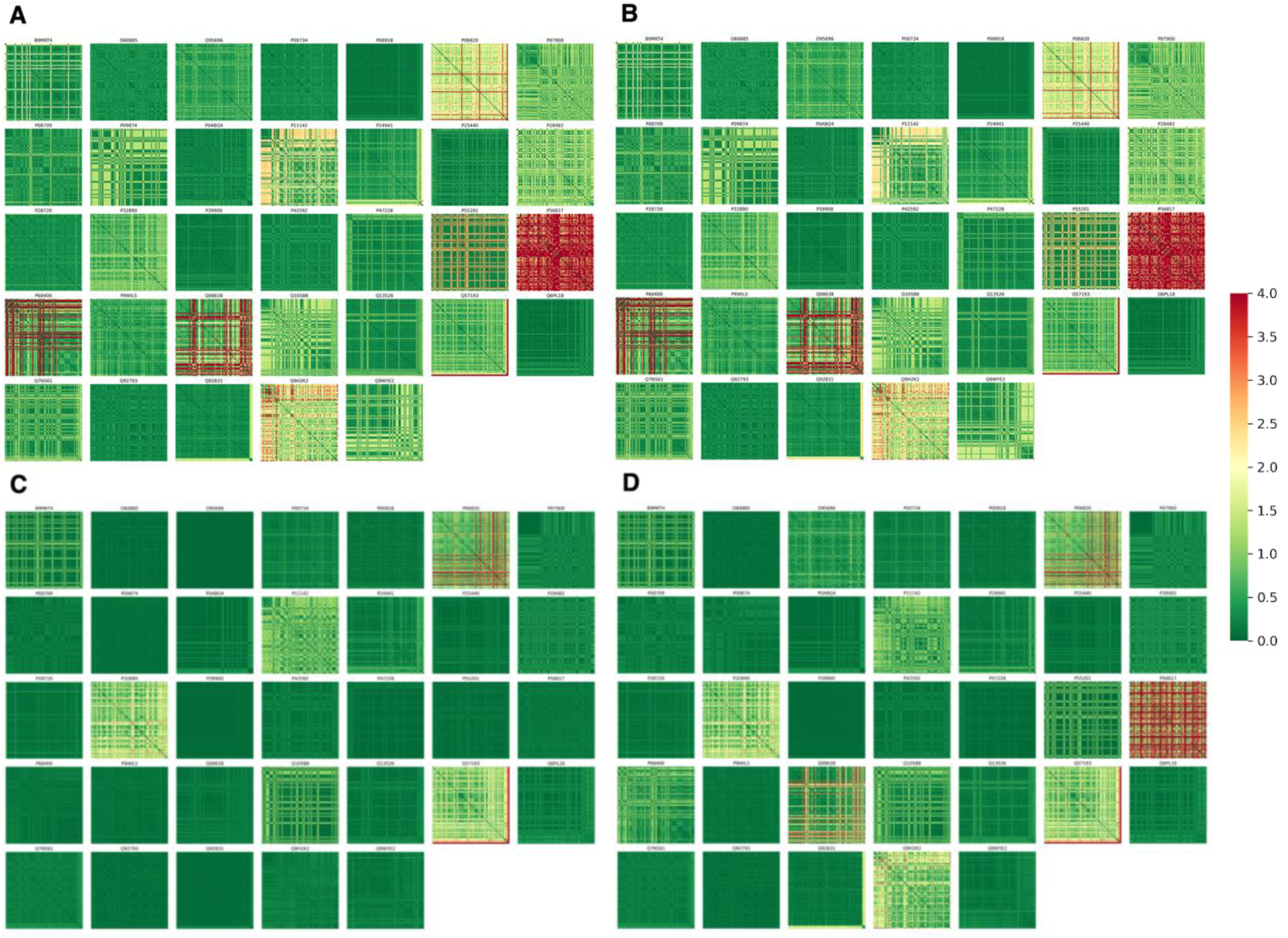
Heatmaps for pairwise RMSDs across the AF ensembles. Shown in (A) whole-protein and top 100 pLDDT ranked models and (B) for the whole chain and best pLDDT per seed, and in (C) for binding site residues for 100 ranked models and in (D) for binding site residues and best pLDDT per seed. The RMSD color scale ranges from 0 Å (dark green) to 4 Å (dark red).

As noted above, we utilized the top 100 model ensembles for ligand docking given the nearly equivalent structural variability of the ensembles generated by each approach and the robust structural prediction accuracy. For these AF ensembles, we also analyzed the impact of the conformational heterogeneity of the AF ensemble on docking outcomes. On average, the AF ensembles were derived from 77.7 unique seeds (**Figure 6**); approximately half of the AF ensembles were derived from 77 or more unique seeds (16/33 ensembles) and the number of seeds represented in the ensemble ranged from 59 to 100. The majority of AF ensembles that yielded a correct (RMSD ≤ 2 Å) or a near-correct (RMSD ≤ 2.5 Å) docking pose in the top 1 or a correct pose in the top 5 were derived from a number of unique seeds at or above the mean (8/10 AF ensembles). All ensembles except for P32890 for which no correct or near correct pose was identified have number of unique seeds below the mean. This indicates that AF ensembles composed of models derived from a greater number of random seeds are more likely to yield favorable docking outcomes. The enhanced performance likely stems from higher heterogeneity in binding site geometries associated with a larger number of random seeds, which raises the probability of sampling binding-optimal states for accommodating the ligand (36).

**FIGURE 6.**
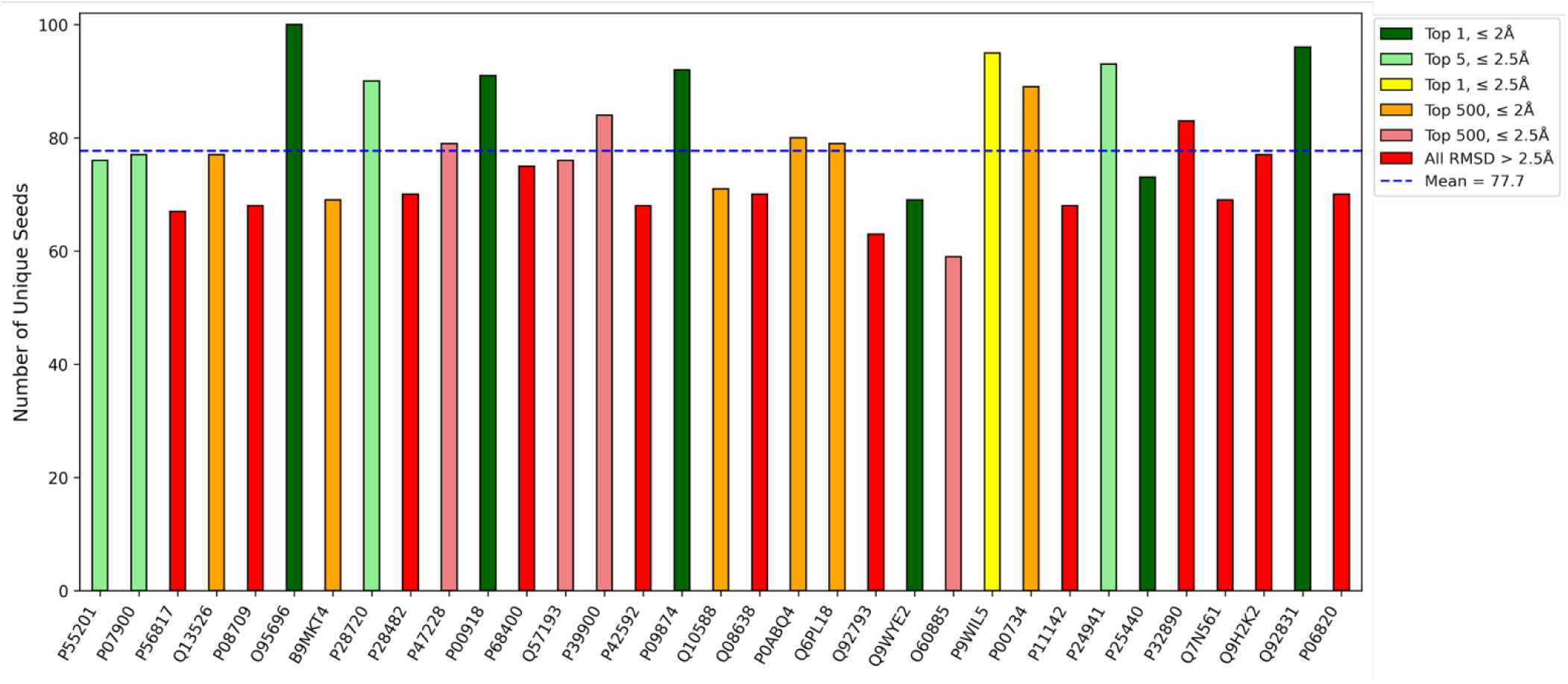
Comparison of docking accuracy vs. number of unique seeds represented in each AF ensemble by protein. Each ensemble consists of top 100 AF models ranked by pLDDT score. Bar colors represent six categories of larger ligand docking outcomes with green representing the most accurate docking and red the least. The mean number of unique seeds per AF ensemble is 77.7.

The distribution of pLDDT scores for the top 100 model AF ensembles and their respective docking outcomes (**Figure 7**) reveals that majority of ensembles show tightly clustered pLDDT scores across both the whole protein (**Fig. 7A**) and the binding site region (**Fig. 7B**), with mean pLDDT values of 93.6 and 95.0, respectively. The whole-protein pLDDTs for 21/33 AF ensembles were at or above the mean, while the remaining 12/33 AF ensembles have pLDDT scores between 87-93. The binding site pLDDTs were somewhat better for 25/33 AF ensembles with scores at or above the mean, while 5 proteins had pLDDTs between 85 and 95, and two of the AF ensemble had poor pLDDT score scores between 70 and 80. Ensembles with favorable and unfavorable docking outcomes are equally distributed above and below the whole-protein pLDDT mean suggesting that high global structural confidence is not tightly correlated to docking success rate (6). However, when looking at the relation to the binding site pLDDTs, the scores of nearly all AF ensembles with correct or near correct docking results (except for P24941 and P28720) lie above the mean. This implies that although high global confidence in protein structure prediction does not guarantee successful ligand docking, the structural accuracy of binding site region may be a more reliable indicator.

**FIGURE 7.**
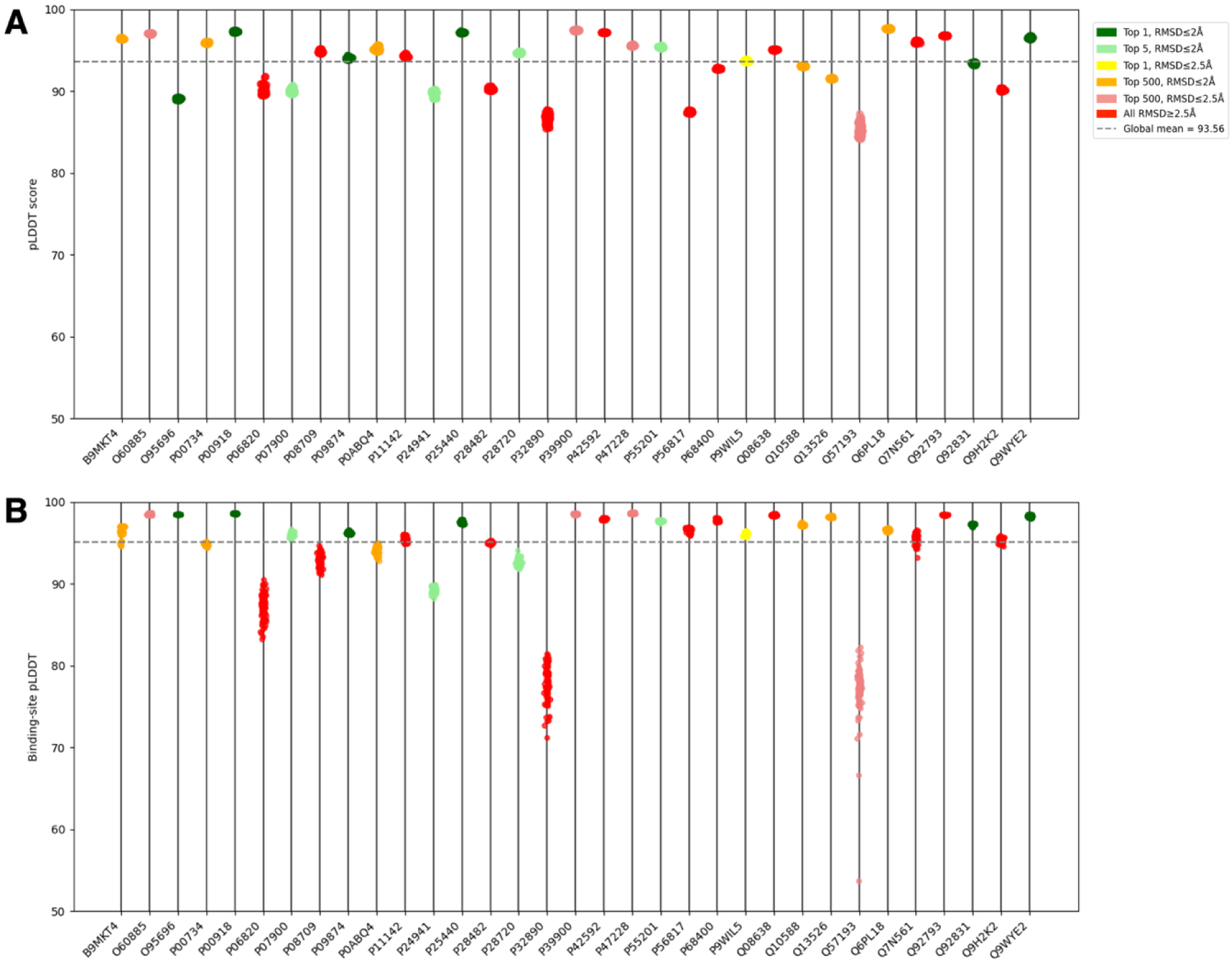
Scatter plots for pLDDT confidence scores. Shown in (A) for the whole-protein and (B) for the binding site regions across the AF ensembles for each protein. Each ensemble consists of top 100 AF models ranked by pLDDT score. Bar colors represent six categories of larger ligand docking outcomes with green representing the most accurate docking and red the least. The mean of the whole-protein and binding site pLDDTs over the test set are 93.6 and 95.0, respectively.

## CONCLUSION

Molecular ligand docking to AF models can perform comparably or even outperform docking to experimentally determined unbound structures. Fragment docking to AF models using unbiased FTMap-identified binding sites yields a greater proportion of correct and near-correct poses in both the top 1 and top 5 ranks. Most importantly, incorporating protein flexibility by docking to multiseed-generated AF ensembles increases the success rate for larger ligands, surpassing results obtained for unbound structures.

These findings introduce a practical workflow for structure-based virtual screening without when no experimental structure is available: (i) utilize FTMap to identify hot spots on an AF model of the protein (without prior knowledge of the native ligand binding site) to assess the ligandability of the protein of interest (34,35,37) and define the docking box, (ii) generate ensembles of AF models using a multiseed approach to account for protein flexibility during docking and (iii) dock ligands to AF ensembles to rapidly screen large ligand libraries to select compounds for experimental testing. This approach addresses the core challenge of limited experimental structure availability and provides an efficient alternative to computationally intensive machine learning methods like Boltz-2 (38) that also can rank ligands.

The quality of binding site representation in AF models remains a critical determinant of docking success. Experimentally determined structures using X-ray crystallography inherently capture the subtleties of protein conformation by taking a real-time “snapshot”. Conversely, AF predicts the protein conformation by leveraging the residue interactions captured by evolutionary relationships in sequence and in all previously determined structures. A limitation of our approach is that AF predictions, while highly accurate globally, may lack the correct small-scale adjustments necessary for binding site integrity. In addition, the proteins in our test set were present in the AF training set (as they were in the other docking studies mentioned (6, 8–10)). While a protein with a completely novel structure may show degradation in the subsequent docking results, capturing conformational variability with multiseed-generated AF ensembles is still expected to mitigate this effect somewhat. Additionally, AF models lack certain biological contexts including complex formation with other macromolecules (e.g., DNA, RNA), the presence of metal ion cofactors, post-translational modifications, and solvation (39). Careful assessment of the impact of such factors on protein conformation with methods like MD, or less computationally intensive methods like normal mode analysis, may be necessary to capture proteins in their native biological state (40, 41).

Several areas for future development of this methodology include: refinement of docking scoring functions specifically for docking to AF models, integration of pLDDT confidence scores; investigation of approaches to integrate more biologically relevant states; exploration of ML-based co-folding predictions as starting points for molecular docking (42–44). The successful use of AF models for unbiased ligand docking highlights the potential for fully integrating deep-learning protein structure prediction approaches into structure-based drug design, thereby helping to streamline the discovery of novel therapeutics for targets without experimentally determined structures.

## Data and Software Availability

All structure data used in this work are from the Protein Data Bank files listed in Table 1. The full test set including bound and unbound structures, AF models, and AF ensembles can be found on the Zenobo repository at https://doi.org/10.5281/zenodo.17289459, along with scripts for analyzing the data. The FTmap server is free for academic and governmental use at https://ftmap.bu.edu/.

## Supporting Information

Figure S1. Visual representation of three different enclosing box configuration for receptor grid generation; Figure S2. Comparison of near correct (RMSD ≤ 2.5 Å) larger ligand poses across the protein test set and induced fit docking for IFD from the Schrodinger release 2024-2 with Glide SP for the initial docking followed Glide XP; Figure S3. Comparison of near correct (RMSD ≤ 2.5 Å) larger ligand poses across the protein test set and induced fit docking for IFD from the Schrodinger release 2025-3 with Glide SP for the initial docking followed Glide SP; Table S2. Assessment of larger ligand docking cases with high RMSD results; Table S3. Fragment docking performance using Glide SP across the test set; Table S3. Larger-ligand docking performance using Glide SP across the test set.

## Author contributions

HM performed computations, analyzed data, wrote manuscript; ML contributed analytic tools and edited manuscript; RB performed computations; RS performed computations; SV oversaw project, edited manuscript, and DJ-M designed research, oversaw project, and edited manuscript.

