## Supplemental Figures S1-S3 and Tables S1-S4. for "ASSESSMENT OF ALPHAFOLD PROTEIN MODELS FOR SMALL-MOLECULE LIGAND DOCKING"

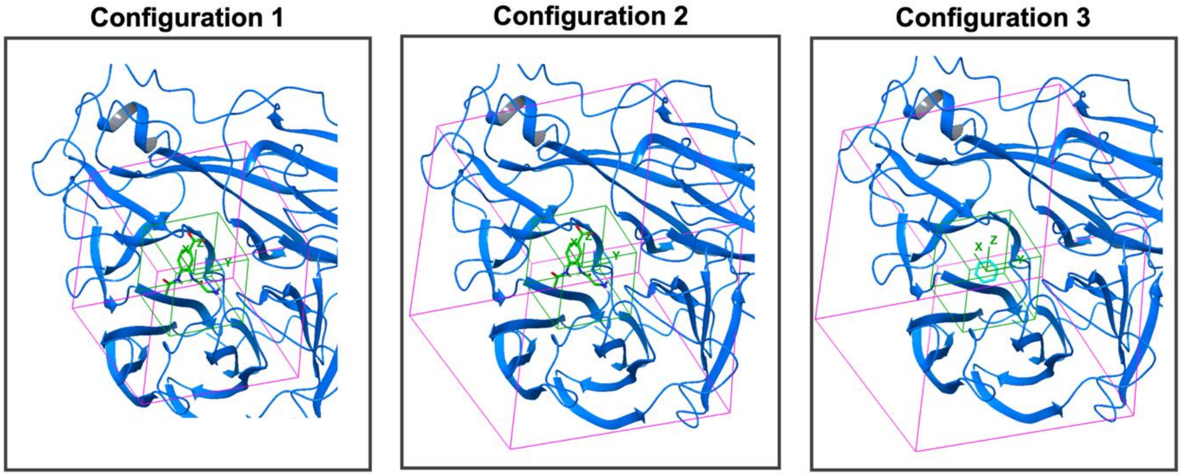


**Figure S1. Visual representation of three different enclosing box configuration for receptor grid generation.** Each panel displays inner and outer (enclosing) boxes that constrain the receptor grid calculations, in green and magenta respectively. Under configuration 1, a bound ligand was used to define box center with outer box size determined based on the size of the ligand. Configuration 2 used the same bound ligand as box center, but the box size was manually adjusted to accommodate ligands up to 20 Å in length. Configuration 3 displays the use of consensus site probe as box center with box size similarly set for ligands ≤ 20 Å.

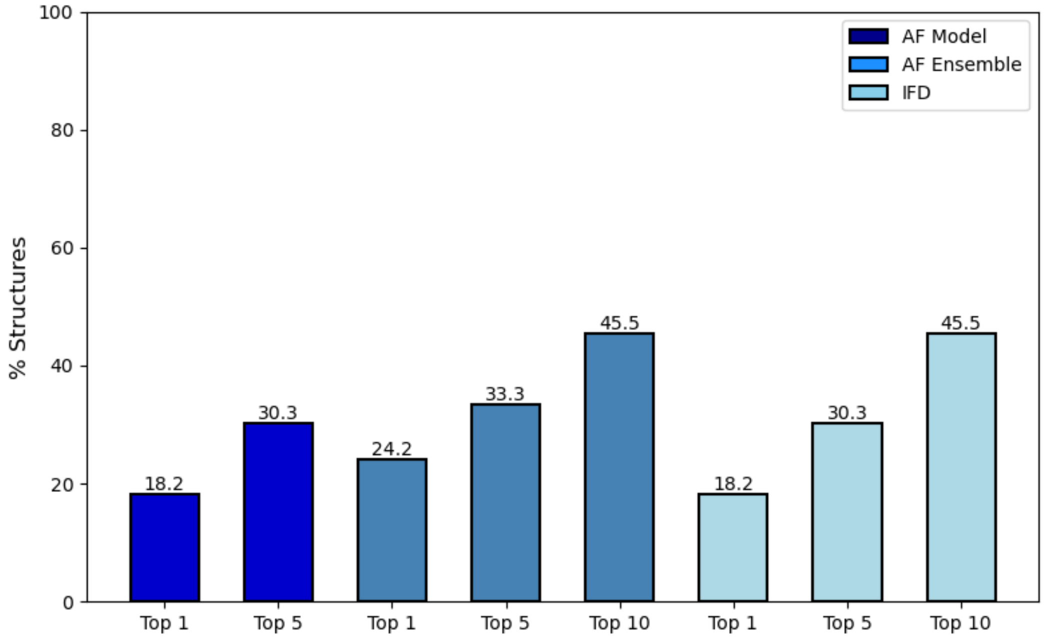


**Figure S2. Comparison of near correct (RMSD ≤ 2.5 Å) larger ligand poses across the protein test set and induced fit docking for IFD from the Schrodinger release 2024-2 with Glide SP for the initial docking followed Glide XP.** Results are shown for box configuration 1 in (A) for the top ranked pose and in (B) for the best-RMSD pose among top 5 ranked poses.

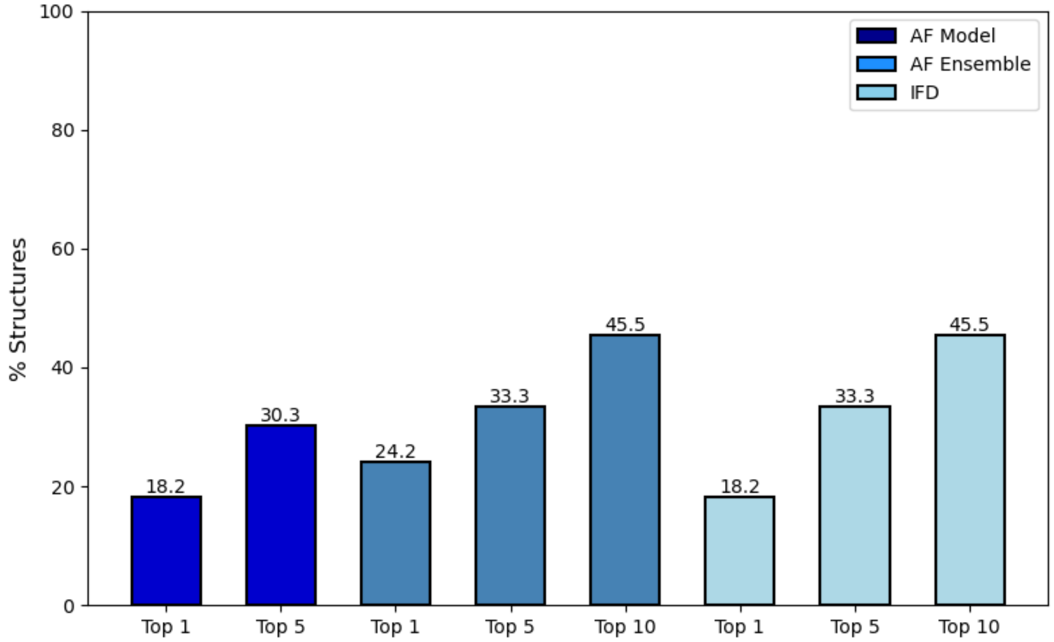


**Figure S3. Comparison of near correct (RMSD ≤ 2.5 Å) larger ligand poses across the protein test set and induced fit docking for IFD from the Schrodinger release 2025-3 with Glide SP for the initial docking followed Glide SP.** Results are shown for box configuration 1 in (A) for the top ranked pose and in (B) for the best-RMSD pose among top 5 ranked poses.

**Table S1. Docking failures due to incorrect binding site identification [a]**

| | | | Fragments | | Larger Ligands | | |
|---|---|---|---|---|---|---|---|
| **UniProt ID** | **Unbound PDB hot spot ≤ 13** | **AF model hot spot ≤ 13** | **Unbound PDB with RMSD ≤ 2.5** | **AF model with RMSD ≤ 2.5** | **Unbound PDB with RMSD ≤ 2.5** | **AF model with RMSD ≤ 2.5** | **AF ensemble with RMSD ≤ 2.5** |
| B9MKT4 | no | no | no | no | no | no | no |
| P68400 | no | yes (Site1) | no | yes | no | no | no |
| P00734 | yes (Site0) | no | no | no | no | no | no |
| Q9H2K2 | yes (Site0) | no | no | no | no | no | no |
| P06820 | no | yes (Site0) | no | yes | no | no | no |

[a] For all other proteins in the test set, FTMap identified a hot spot within 2.5 A of either the fragment or larger ligand.

**Table S2. Assessment of larger ligand docking cases with high RMSD results [a]**

| UniProt ID | Unbound RMSD [Å] | Explanation | AF RMSD [Å] | Explanation |
|---|---|---|---|---|
| B9MKT4 | 29.7 | The binding site is identified only with low-ranking probe clusters: CS02 (12 probes), CS06 (6 probes). Possibly due to 3 Ca2+ ions present in the binding site of the bound structure (1) | 14.7 | The binding site mapped with CS00 (13 probes) but it is 3.6 Å away and the larger ligand is a disaccharide with 62 tautomeric states |
| P68400 | 14.8 | The binding site is identified with CS01(18 probes), but it is 4.3 Å away from the ligand tail, and with CS04 (9 probes) and CS07 (4 probes) that do overlap with the ligands | 13.4 | The binding site is identified with CS01 (17 probes) and overlaps with both ligands. The larger ligand has 11 rotatable bonds |
| P39900 | 14.5 | The binding site is identified only with low-ranking CS03 (10 probes) that overlaps with the ligands. There is a catalytic Zn ion present in the binding site of the bound structures (2) | 7.0 | The binding site is identified with CS00 (26 probes). Docked ligand poses are incorrectly oriented. There is a catalytic Zn ion in the binding the bound structures that is absent in the calculations |
| P32890 | 9.6 | The binding site is correctly identified with CS02 (15 probes) which overlaps with both ligands. Docked poses are incorrectly oriented possibly due the large ligand having 9 rotatable bonds | 16.0 | The binding site is identified with CS03 (14 probes) which overlaps with both ligands. Docked poses are also incorrectly oriented but more removed from the bound reference compared to poses in the unbound structure |
| Q9H2K2 | 7.6 | The binding site is identified with CS00 (19 probes), CS01 (18 probes) and CS02 (17 probes). Part of the ligand is correctly oriented. The larger ligand has 6 rotatable bonds | 22.6 | There are no consensus sites in the binding site |

[a] For unbound protein structures and AF models of five proteins, docked poses of larger ligands yielded disproportionally high RMSD values relative to the bound reference. This table summarizes the  rationale for these outliers alongside their corresponding RMSD values.

**Table S3. Fragment docking performance using Glide SP across the test set**

| | | RMSD ≤ 2 [Å] | | | RMSD ≤ 2.5 [Å] | | |
|---|---|---|---|---|---|---|---|
| **Box Conf.** | **Pose Rank[a]** | **X-ray Bound [%]** | **X-ray Unbd. [%]** | **AF Models [%]** | **X-ray Bound [%]** | **X-ray Unbd. [%]** | **AF Models [%]** |
| 1 | Top 1 | 15.2 | 15.2 | 24.2 | 30.3 | 18.2 | 33.3 |
| | Top 5 | 60.6 | 45.5 | 45.5 | 66.7 | 48.5 | 54.5 |
| 2 | Top 1 | 18.2 | 18.2 | 27.3 | 30.3 | 18.2 | 39.4 |
| | Top 5 | 54.5 | 51.5 | 45.5 | 63.6 | 45.5 | 54.5 |
| 3 | Top 1 | 24.2 | 21.2 | 21.2 | 33.3 | 27.3 | 33.3 |
| | Top 5 | 54.5 | 45.5 | 36.4 | 57.6 | 57.6 | 51.5 |

[a] Results are for the top ranked pose and the lowest RMSD pose among top 5 poses

**Table S4. Larger-ligand docking performance using Glide SP across the test set**

| | | RMSD ≤ 2 [Å] | | | | RMSD ≤ 2.5 [Å] | | | |
|---|---|---|---|---|---|---|---|---|---|
| **Box Conf.** | **Pose Rank[a]** | **X-ray Bound [%]** | **X-ray Unbd. [%]** | **AF Models [%]** | **AF Ens. [%]** | **X-ray Bound [%]** | **X-ray Unbd. [%]** | **AF Models [%]** | **AF Ens. [%]** |
| 1 | Top 1 | 24.2 | 18.2 | 15.2 | 12.2 | 30.3 | 21.2 | 15.2 | 15.2 |
| | Top 5 | 36.4 | 30.3 | 27.3 | 21.2 | 48.5 | 30.3 | 30.3 | 27.3 |

[a] Results are for the top ranked pose and the lowest RMSD pose among top 5 poses
